# Tissue-specific Genome Editing *in vivo* by MicroRNA-repressible Anti-CRISPR Proteins

**DOI:** 10.1101/631689

**Authors:** Jooyoung Lee, Haiwei Mou, Raed Ibraheim, Shun-Qing Liang, Wen Xue, Erik Sontheimer

**Affiliations:** RNA Therapeutics Institute, University of Massachusetts Medical School, Worcester, Massachusetts, U.S.A.; Program in Molecular Medicine, University of Massachusetts Medical School, Worcester, Massachusetts, U.S.A.

**Keywords:** Cas9, anti-CRISPR, microRNA, tissue-specific editing

## Abstract

CRISPR-Cas systems are bacterial adaptive immune pathways that have revolutionized biotechnology and biomedical applications. Despite the potential for human therapeutic development, there are many hurdles that must be overcome before its use in clinical settings. Some clinical safety concerns arise from persistent activity of Cas9 after the desired editing is complete, or from editing activity in unintended cell types or tissues upon *in vivo* delivery [e.g. by adeno-associated viruses (AAV)]. Although tissue-specific promoters and serotypes with tissue tropisms can be used, suitably compact promoters are not always available for desired cell types, and AAV tissue tropisms are not absolute. To reinforce tissue-specific editing, we exploited anti-CRISPR proteins (Acrs), which are proteins evolved as countermeasures against CRISPR immunity. To inhibit Cas9 in all ancillary tissues without compromising editing in the target tissue, we established a flexible platform in which an *Acr* transgene is repressed by endogenous, tissue-specific microRNAs (miRNAs). We demonstrate that miRNAs regulate the expression of an *Acr* transgene bearing miRNA-binding sites in its 3’ UTR, and control subsequent genome editing outcomes in a cell-type specific manner. We also show that the strategy is applicable to multiple Cas9 orthologs and their respective Acrs. Furthermore, we demonstrate that *in vivo* delivery of Cas9 and Acrs that are targeted for repression by liver-specific miR-122 allow editing in the liver while Acrs devoid of miRNA regulation prevent Cas9 activity. This strategy provides additional safeguards against off-tissue genome editing by confining Cas9 activity to selected cell types.

## Introduction

Clustered, regularly interspaced, short, palindromic repeats (CRISPR) and CRISPR-associated (*cas*) genes comprise prokaryotic adaptive immune defense systems that are classified into two major classes and multiple types and subtypes (e.g. II-A, -B, and -C) (Makarova et al. 2018). Cas9s are monomeric effector proteins in type II systems that can target nearly any DNA sequence when guided by a CRISPR RNA (crRNA) base paired with a trans-activating RNA (tracrRNA), or as a fused form of both RNAs known as single guide RNA (sgRNA) (Deltcheva et al. 2011; Garneau et al. 2010; Jinek et al. 2012). The robustness and ease of Cas9 programmability have greatly facilitated its rapid adoption in genome editing and modulation (Komor et al. 2017). As medical, agricultural, and environmental technologies advance, safety concerns must be considered and addressed, especially with potential human therapeutics. *In vivo* therapeutics will often require not only precise editing at the intended genomic site but also in the intended tissue, given the possible risks of unwanted double-strand break (DSB) induction. For example, Cas9-induced DSBs can elicit translocations that can be associated with heritable disorders or various kinds of cancer, or large deletions and other rearrangements (Jiang et al. 2016; Maddalo et al. 2014; Kosicki et al. 2018). Moreover, some delivery modalities such as viral vectors are likely to affect many cell types and tissues beyond the intended therapeutic target (Hinderer et al. 2018). AAV is currently the most widely used transgene delivery vector for therapeutic applications in preclinical and clinical settings. Different AAV serotypes have some tissue tropism, however, they can still infect broad ranges of tissues *in vivo* (Gao et al. 2004). Although tissue-specific promoters can be used to drive transgene expression in particular cell types (Walther and Stein 1996), some target tissues lack promoters that are sufficiently active, specific, or small for AAV deployment. These limitations necessitate the development of new regulatory strategies to enforce tissue specificity for *in vivo* applications.

Although several means of regulating genome editing activities have been reported, a prominent recent advance has resulted from the discovery of anti-CRISPR (Acr) proteins (Bondy-Denomy et al. 2013). Acrs are small proteins encoded by bacteriophages and other mobile genetic elements that have evolved as natural countermeasures against CRISPR-Cas immunity. Type II Acrs targeting Cas9 orthologs (Pawluk et al. 2016; Rauch et al. 2017; Hynes et al. 2017, 2018), as well as the recently-discovered type V Acrs targeting Cas12a (Watters et al. 2018; Marino et al. 2018), are of particular interest because they can potentially provide temporal, spatial, or conditional control over established genome editing systems. Applications of Acrs have been demonstrated in bacteria (Marshall et al. 2018; Rauch et al. 2017), in yeasts to inhibit gene drives (Goeckel et al. 2019), and in mammalian cells to modulate genome editing, dCas9-based imaging, epigenetic modification, and genetic circuits (Pawluk et al. 2016; Rauch et al. 2017; Shin et al. 2017; Bubeck et al. 2018; Liu et al. 2018; Nakamura et al. 2019).

To improve current technologies that regulate the tissue specificity of editing, we have developed an Acr-based approach to inhibit Cas9 in all ancillary tissues while allowing editing in the target tissue. To spatially regulate Acr expression, we exploited endogenous tissue-specific microRNAs (miRNAs) to repress Acr expression in the target tissue. MiRNAs are a class of small regulatory RNAs whose mechanisms of messenger RNA (mRNA) regulation are extensively studied (Jonas and Izaurralde 2015). These RNAs load into an argonaute protein (e.g. Ago2) to form RNA-induced silencing complexes (RISCs) that recognize complementary sequences present in mRNA targets, leading to translational repression and mRNA destabilization (Bartel 2018). In mammalian cells, Ago2-loaded miRNAs can subject extensively or perfectly complementary mRNA targets to endonucleolytic cleavage, enabling strong downregulation. Since miRNA response elements (MREs) are very small (∼22 nucleotides or less), this regulatory modality places minimal burden on AAV vector capacity, which is limited to ∼4.8 kb. Moreover, large numbers of mammalian cell and tissue types express specific combinations of tissue-restricted miRNAs (Lagos-Quintana et al. 2002).

Here we establish a flexible platform in which an *Acr* transgene is repressed by endogenous, tissue-specific miRNAs to control Acr expression spatially. We demonstrate that miRNAs can regulate the expression of an *Acr* transgene bearing miRNA-binding sites in its 3’ untranslated region (UTR) and control subsequent genome editing outcomes in a cell-type specific manner. We also show that the strategy is applicable to multiple Cas9 orthologs and their respective Acrs, including the widely-used *Streptococcus pyogenes* (SpyCas9) (Cong et al. 2013; Mali et al. 2013; Jinek et al. 2013; Cho et al. 2013; Hwang et al. 2013) as well as the more readily AAV-deliverable Cas9 orthologs from *Neisseria meningitidis* (Nme1Cas9 and Nme2Cas9) (Ibraheim et al. 2018; Edraki et al. 2018). Furthermore, we have expressed anti-CRISPR proteins in mice to achieve efficient inhibition of Cas9-mediated genome editing *in vivo* without detectable toxicity. We show that co-delivery of Cas9, guide RNA, and miR-122-repressible Acr transgenes allow editing in the liver (the only tissue where miR-122 is expressed), while an otherwise identical Acr transgene that lacks any miR-122 MREs effectively prevent Cas9 activity. This strategy establishes the *in vivo* efficacy of Acrs in mammals and provides the basis for restriction of undesired off-tissue editing by confining Cas9 activity to selected cell types.

## Results

### AAV delivery of all-in-one Nme1Cas9/sgRNA results in editing in various tissues

Previously, our group has used all-in-one AAV8 to deliver a human-codon-optimized Nme1Cas9 for genome editing *in vivo* (Ibraheim et al. 2018). Nme1Cas9 is smaller and less prone to off-target editing than the widely used SpyCas9 (Amrani et al. 2018). Upon delivery of all-in-one rAAV8 viruses expressing hNme1Cas9 driven by a ubiquitous U1a promoter and sgRNA via tail vein injection, we observed high editing efficiency in liver tissues collected 50 days post-injection (Ibraheim et al. 2018). To gauge editing efficiencies in non-target tissues outside of the liver, tissues from cardiac and skeletal muscle (gastrocnemius muscle) as well as kidney and brain were collected and analyzed (Supplemental Fig. 1). Although lower than the editing observed in liver tissues (51.33±4.93 %), appreciable indel frequencies were observed in different organs, especially in the heart (22.33±3.79 %) (Supplemental Fig. 1). This is consistent with previous reports that AAV8 effectively transduces mouse hepatocytes but also infects skeletal and cardiac muscles (Nakai et al. 2005) as well as brain at high doses (Zincarelli et al. 2008). These observations, along with the known multi-tissue tropisms of other AAV serotypes (Zincarelli et al. 2008), underscore the potential benefit of using miRNA-repressible *Acr* transgenes to reinforce tissue-specific editing.

### A strategy for microRNA-regulated anti-CRISPR proteins

Endogenous miRNA-mediated post-transcriptional gene silencing has proven to be an effective and tissue-specific approach to regulate transgene expression upon AAV delivery *in vivo* (Xie et al. 2011). Delivery of Cas9/sgRNA via AAV has the potential to induce editing in multiple transduced tissues (e.g. heart, skeletal muscles etc.); however, co-delivery of the miRNA-repressible Acr will inhibit editing in such non-target tissues due to the latter’s lack of tissue-specific miRNAs (and therefore their inability to silence the expression of the Acr inhibitor). In the case of the liver-specific miRNA miR-122, in the target tissue the *Acr* gene with miR-122 MREs will be repressed, enabling Cas9-mediated editing (Fig. 1A). In contrast, off-tissue editing (e.g. in cardiac and skeletal muscle, Supplemental Fig. 1) will be inhibited by the Acr, since those extrahepatic tissues lack miR-122 and therefore fail to silence Acr expression. To validate this concept, we chose two well-established Cas9-Acr combinations: AcrIIC3_*𝒩me*_ and Nme1Cas9/Nme2Cas9 (Type II-C; (Pawluk et al. 2016; Edraki et al. 2018)) as well as AcrIIA4_*Lmo*_ and SpyCas9 (Type II-A; (Rauch et al. 2017)). Nme2Cas9 is a recently reported Cas9 ortholog that has a dinucleotide (N_4_CC) protospacer adjacent motif (PAM) (Edraki et al. 2018), enabling a target site density comparable to that of SpyCas9 (NGG PAM). A type II-C Nme1Cas9/Nme2Cas9 inhibitor, AcrIIC3_*𝒩me*_, limits target DNA affinity (Harrington et al. 2017; Zhu et al. 2019). AcrIIA4_*Lmo*_ inhibits the widely-used SpyCas9 and also prevents DNA binding, in this case by occluding the PAM-binding cleft (Rauch et al. 2017; Dong et al. 2017; Shin et al. 2017; Yang and Patel 2017). For our *in vitro* validations, both Cas9 and Acr expression vectors were driven by the cytomegalovirus (CMV) promoter. We generated codon-optimized Acr expression vectors identical in every respect except for the presence or absence of MREs in the 3’ UTR (Supplemental Table 1). Since miR-122 is a well-validated miRNA that is highly expressed specifically in hepatic cells, we decided to validate the concept using this miRNA. We placed three tandem miR-122 binding sites (3xmiR122BS) in the 3’ UTR of each *Acr* gene, which also included a C-terminal mCherry fusion to enable expression to be detected by fluorescence microscopy or flow cytometry (Fig. 1B). Fusion of heterologous domains do not compromise the inhibitory potency of these Acrs (Goeckel et al. 2019; Nakamura et al. 2019).

**Figure 1.**
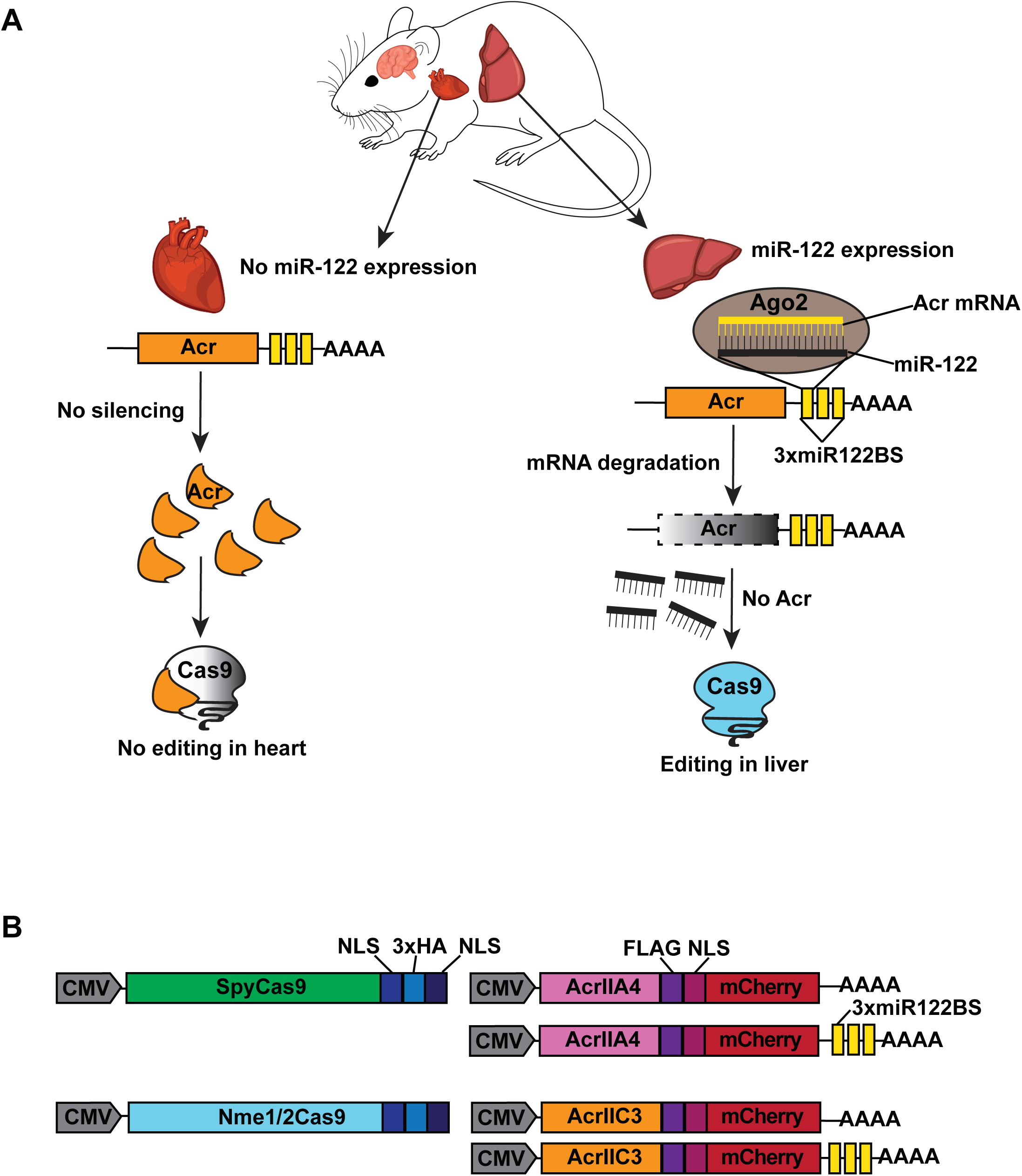
Overview of Cas9 and microRNA-repressible anti-CRISPR system. A. MiRNA-repressible anti-CRISPR and Cas9 editing strategy as designed for use in mice. As an example, miR-122 can be used to achieve liver-specific editing. Upon systemic delivery of Cas9 *in vivo* (e.g. via viral vectors), tissues receiving Cas9 and sgRNA potentially result in genome editing; however, co-delivery of miRNA-repressible anti-CRISPR proteins will prevent such editing in non-target tissues that lack miR-122, as depicted in the heart (left). In liver, anti-CRISPR transcripts with perfectly complementary miR-122 binding sites will undergo Ago2-mediated mRNA degradation, and the resulting silencing of the Acr will permit Cas9 editing in the liver (right). B. A schematic of expression vectors for Cas9 orthologs from type II-A (SpyCas9) and II-C (Nme1Cas9 and Nme2Cas9) systems, along with their respective anti-CRISPR proteins, AcrIIA4_*Lmo*_ and AcrIIC3_*𝒩me*_. The Acr expression constructions were generated with or without three tandem, perfect complementary miRNA-122 binding sites in the 3’ UTR. CMV, cytomegalovirus promoter; NLS, nuclear localization signal; AAAA, poly-A tail.

### Validation of microRNA-repressible anti-CRISPR expression vectors

We used a human hepatocellular carcinoma cell line (Huh-7) that abundantly expresses miR-122, in contrast to non-hepatic cell lines such as human embryonic kidney (HEK293T) cells (Fukuhara et al. 2012). As an initial test of miR-122 repression of Acr expression, we transfected cells with plasmids expressing AcrIIC3-FLAG-mCherry-3xmiR122BS, AcrIIA4-FLAG-mCherry-3xmiR122BS, or their respective control vectors lacking the miR-122 binding sites (Fig. 1B). A separate GFP expression plasmid was also included to indicate transfection efficiencies in each cell line. When these vectors were transiently transfected, the expression of mCherry-fused Acr with miR-122 MREs was dramatically suppressed in Huh7 cells whereas Acr-mCherry lacking 3xmiR122BS was still well expressed (Fig. 2A). In HEK293T cells, there was no discernible difference in mCherry signal from the Acr and Acr-3xmiR122BS constructs based on both fluorescence microscopy and flow cytometry (Fig. 2B). Acr expression was also confirmed by anti-FLAG western blot analysis (Fig. 2). Compared to HEK293T cells, transfection efficiency was lower in Huh-7 cells as indicated by a decrease in overall GFP and mCherry signals (Fig. 2A). Nevertheless, fluorescence microscopy, flow cytometry, and Western blot analysis consistently revealed effective reductions of both AcrIIC3-3xmiR122BS and AcrIIA4-3xmiR122BS expression in Huh-7, but not in HEK293T cells. Expression of Acrs lacking miR-122 MREs was unaffected in both cell lines, consistent with effective regulation of Acr by miR-122 only in hepatic cells.

**Figure 2.**
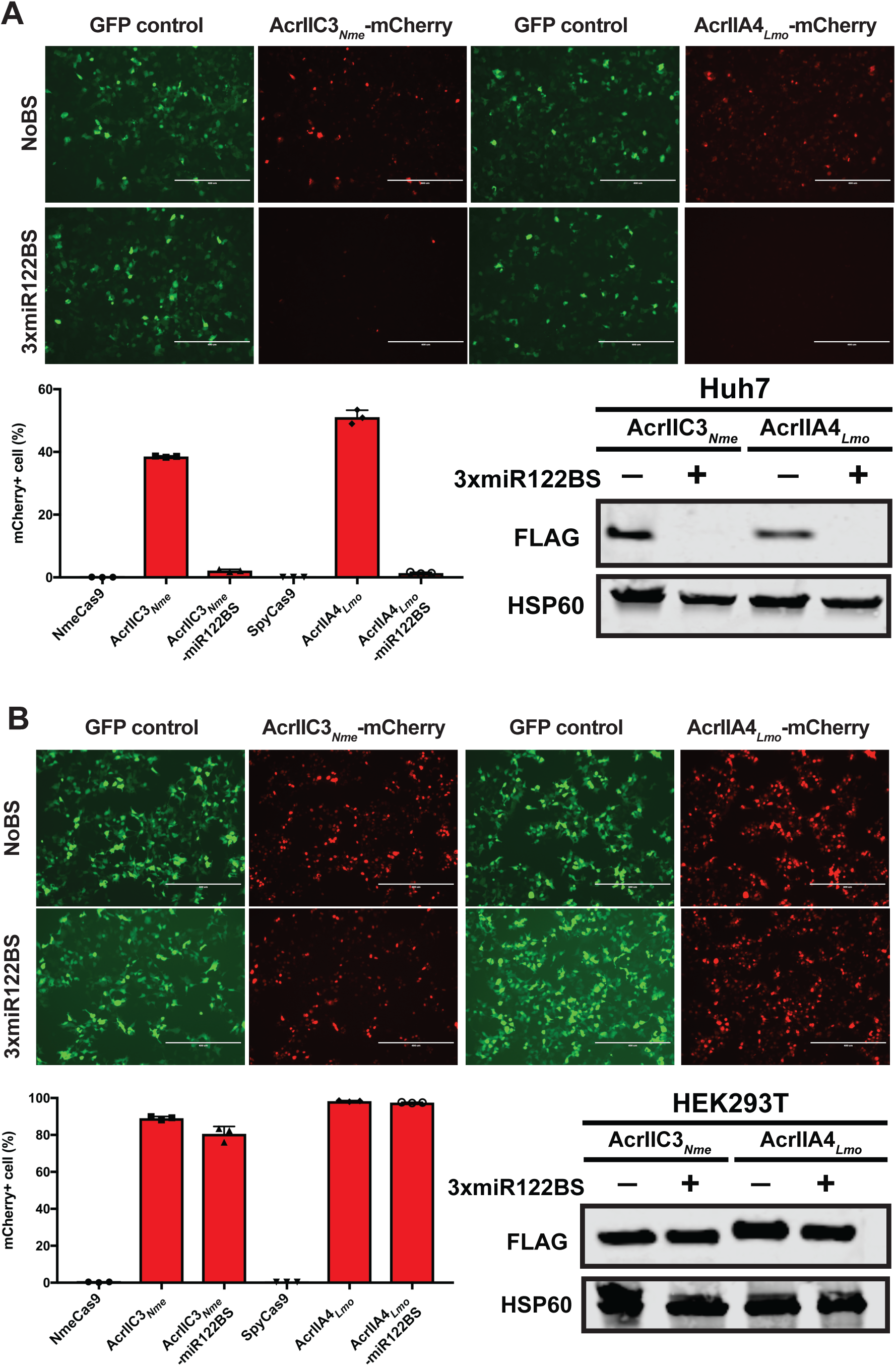
Validation of miRNA regulation of anti-CRISPR expression in cultured cells. (A, B) Hepatocyte-specific silencing of anti-CRISPR expression. Plasmid vectors shown in Fig. 1B encoding either AcrIIC3_*𝒩me*_-mCherry or AcrIIA4_*Lmo*_-mCherry, with or without miR-122 MREs, were transfected into (A) human hepatoma (Huh7) cells or (B) non-hepatic HEK293; only the former express miR-122. The expression of mCherry and GFP was visualized by fluorescence microscopy (top) and analyzed by flow cytometry (bottom left). The percentage of mCherry-positive cells in each transfection was normalized to transfection of the control GFP-expressing plasmid. Anti-CRISPR protein expression was also confirmed by western blot against the 1xFLAG epitope (bottom right). Heat shock protein 60 (HSP60) was used as a loading control. Scale bar, 400 µm.

### MicroRNA repression enables escape from anti-CRISPR inhibition during genome editing in hepatocytes

Having demonstrated that anti-CRISPR repression in hepatocyte-derived cells can be conferred by miR-122 MREs, we then tested whether this repression is sufficient to allow genome editing by Cas9 orthologs (SpyCas9, Nme1Cas9 and Nme2Cas9). We transiently transfected separate expression plasmids for Cas9, a cognate sgRNA, and an Acr, with the latter construct either including or omitting miR-122 binding sites. We chose validated, endogenous sites in the human genome for each Cas9 ortholog (Fig. 3): the Nme1Cas9 target site NTS33 in the *VEGFA* gene (Fig. 3A), the Nme2Cas9 target site TS6 in the *LINC01588* gene (Fig. 3B), and the SpyCas9 target site 1617 in the *BCL11A* enhancer (Fig. 3C) (Amrani et al. 2018; Edraki et al. 2018; Wu et al. 2019). In HEK293T cells, AcrIIC3_*𝒩me*_ and AcrIIA4_*Lmo*_ robustly inhibited genome editing by Nme1/2Cas9 and SpyCas9, respectively, as expected (Pawluk et al. 2016; Rauch et al. 2017) (Fig. 3). The presence or absence of miR-122 MREs had no significant effect on editing inhibition in this non-miR-122-expressing cell type. Although the editing efficiency was variable among Cas9 orthologs at these target sites, and although transfection efficiencies were reduced in Huh-7 cells, AcrIIC3_*𝒩me*_ and AcrIIA4_*Lmo*_ also prevented editing in this cell type when expressed from constructs that lack miR-122 MREs. By contrast, Acrs plasmids that incorporated miR-122 MREs in the 3’UTRs failed to inhibit Cas9 editing in Huh-7 cells, as indicated by editing efficiencies that were similar to the no-Acr control (Fig. 3). This trend was true for all three Cas9 orthologs tested.

**Figure 3.**
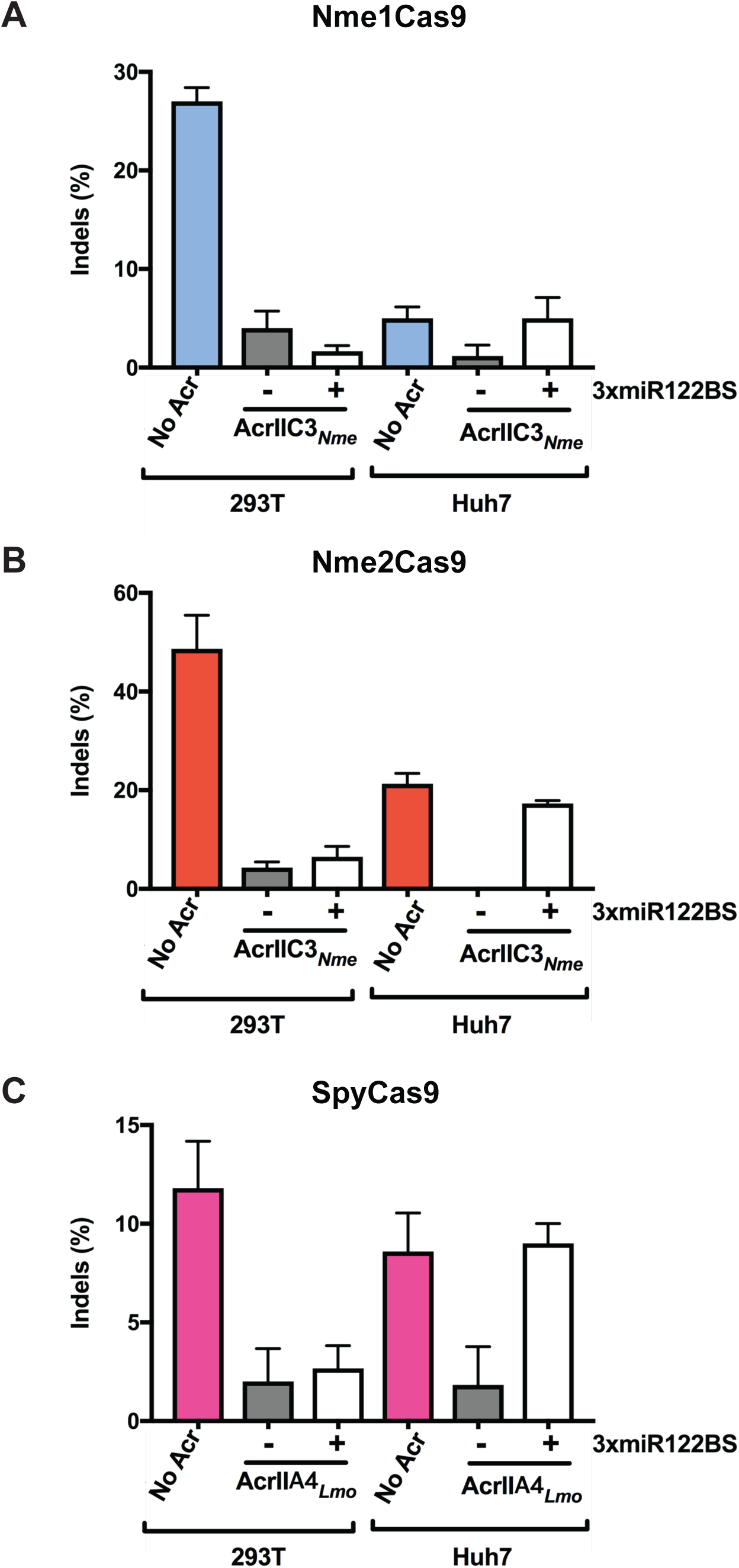
Hepatocyte-specific genome editing by Nme1Cas9, Nme2Cas9 and SpyCas9 in cultured cells. (A-C) HEK293T and Huh7 cells were transiently transfected with plasmids encoding (A) Nme1Cas9 and an sgRNA targeting the *VEGFA* locus, (B) Nme2Cas9 and an sgRNA targeting *LINC01588*, and (C) SpyCas9 and an sgRNA targeting the *BCL11A* enhancer. (A, B) AcrIIC3_*𝒩me*_ constructs with or without 3xmiR122BS were co-transfected with the Cas9 and sgRNA constructs as indicated. (C) AcrIIA4_*Lmo*_ with or without 3xmiR122BS were co-transfected with SpyCas9 and its sgRNA. Data represent mean ± s.e.m with at least 3 replicates. Editing efficiencies are measured by TIDE.

### MiR-122-dependent *in vivo* genome editing conferred by an anti-CRISPR protein

For our *in vivo* tests we focused on Nme2Cas9, due to its compact size, high target site density, and relative lack of off-target editing, all of which are advantageous for therapeutic development. We used a previously validated all-in-one AAV vector that expresses Nme2Cas9 from the minimal U1a promoter, as well as a U6 promoter-driven sgRNA targeting *Rosa26* (Ibraheim et al. 2018; Edraki et al. 2018) (Fig. 4A). We also generated AcrIIC3_*𝒩me*_ expression plasmids driven by the strong CB-PI promoter and associated expression elements; in addition, these AcrIIC3_*𝒩me*_ constructs either included or omitted the three tandem miR-122 MREs in the 3’ UTR (Fig. 4A). For *in vivo* delivery we used hydrodynamic injection, which is a non-viral method of transient hepatocyte transfection that allows expression from naked DNA plasmids (Zhang et al. 1999). This injection method delivers DNA to ∼20% of hepatocytes for transient expression and has minimal transgene expression in organs other than the liver. Since miR-122 is abundant in the liver, and because Cas9 delivered to the liver by hydrodynamic injection can induce editing (Xue et al. 2014), this experimental approach enables tests of liver-specific editing (and inhibition of editing) in the presence or absence of Acr expression. Plasmids were injected into adult, wild-type C57BL/6 mice via tail vein and liver tissues were collected at 7 days post-injection (Fig. 4B). To determine the effective dose of Acr plasmid needed to inhibit Nme2Cas9 editing *in vivo*, we co-injected varying Cas9:Acr plasmid ratios (1:1, 1:1.5, and 1:2). AcrIIC3_*𝒩me*_ efficiently inhibited Nme2Cas9 editing at all ratios tested (Fig. 4C). No apparent liver damage was detected in the liver tissues following staining with haemotoxylin and eosin (H&E) (Supplemental Fig. 2). Once we defined the necessary plasmid dose, we subjected three groups of mice to hydrodynamic injection with plasmid combinations that included Nme2Cas9 with (*i*) no Acr, (*ii*) AcrIIC3_*𝒩me*_, and (*iii*) AcrIIC3_*𝒩me*_-3xmiR122BS (Fig. 4A). In the livers of mice receiving no Acr, Nme2Cas9 yielded a mean editing efficiency of 4.2±0.6% (n = 6 mice), similar to levels seen previously with this and other Cas9 orthologs upon hydrodynamic injection (Ibraheim et al. 2018; Xue et al. 2014). As expected, co-injection of AcrIIC3_*𝒩me*_ plasmid strongly reduced the editing efficiency to 1±0.5% (*P* = 0.0025). By contrast, AcrIIC3_*𝒩me*_-3xmiR122BS failed to inhibit Nme2Cas9 editing, with the indel efficiency comparable to no Acr group (6.7±1.1%, Fig. 4D). We confirmed the expression of Nme2Cas9 in all three groups by immunohistochemistry (IHC) against the 3xHA epitope (Supplemental Fig. 3). We were unable to detect AcrIIC3_*𝒩me*_ by IHC against the FLAG epitope in mice injected with AcrIIC3_*𝒩me.*_ It is possible that 1xFLAG tag is too weak for IHC detection. However, we ruled out the possibility of injection failures by including control plasmids in our experiment. Specifically, we co-injected additional plasmids encoding a Sleeping Beauty transposon system (Ivics et al. 1997) that integrates an mCherry expression cassette into the mouse genome to report on the success of plasmid injection. In all three groups of injected mice, we observed mCherry expression in liver tissue sections from injected mice by IHC (Supplemental Fig. 3), confirming successful liver transfection. In summary, consistent with our results in human Huh-7 cells, endogenous miR-122 in mouse hepatocytes *in vivo* can be exploited to repress Acr expression, and therefore allow tissue-specific Cas9 genome editing, in liver tissues.

**Figure 4.**
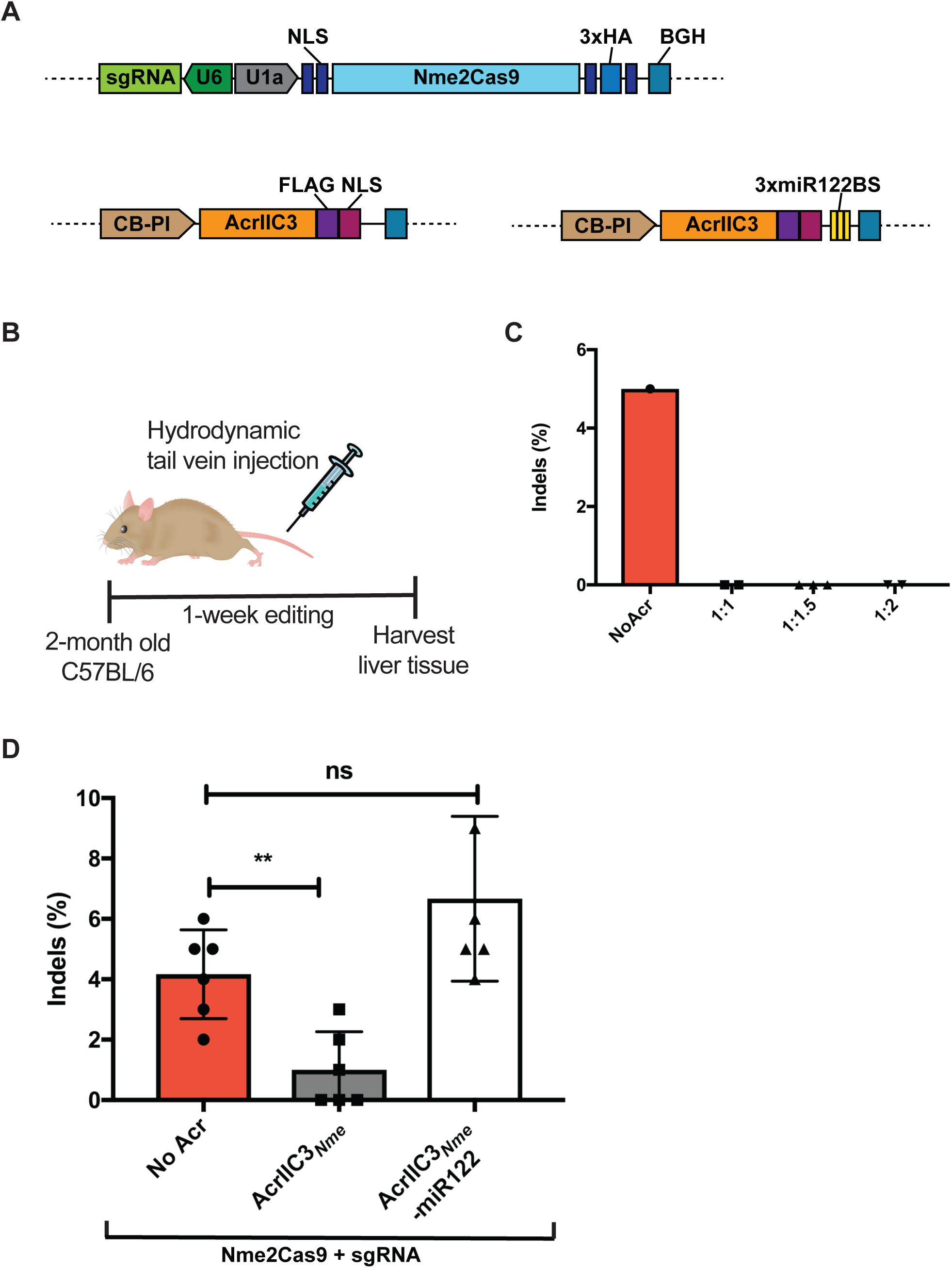
Acr inhibition of Nme2Cas9 editing *in vivo*, and release from inhibition by the liver-specific miRNA, miR-122. A. Plasmids used for *in vivo* studies to drive the expression of Nme2Cas9/sgRNA and AcrIIC3_*𝒩me*_, respectively. U1a, murine promoter; BGH, bovine growth hormone polyA signal; CB-PI, cytomegalovirus-enhancer, chicken β-actin (CB) promoter with SV40-derived mini-intron. B. A schematic of mouse studies. Plasmid vectors shown in (A) are administered into 8-to 10-week-old C56BL/6 mice by hydrodynamic tail vein injection. Liver tissues were collected one week after injection. C. Dose titration of Nme2Cas9/sgRNA plasmid to AcrIIC3_*𝒩me*_ plasmid *in vivo*. Percentage of indels at the *Rosa26* target in the livers of C57Bl/6 mice measured by TIDE after hydrodynamic injection of Nme2Cas9/sgRNA and AcrIIC3_*𝒩me*_ plasmids at mass ratios of 1:1, 1:1.5, and 1:2. D. Genome editing in the liver by Nme2Cas9 is inhibited by AcrIIC3_*𝒩me*_ but restored when AcrIIC3_*𝒩me*_-3xmiR122BS is silenced. Indel percentages at the *Rosa26* locus in the livers of C57Bl/6 mice was measured by TIDE after hydrodynamic injection of Nme2Cas9/sgRNA plasmid, along with anti-CRISPR plasmids with or without 3xmiR122BS. N = 6 mice per group. ns = not significant, p < 0.01 by unpaired, two-tailed t-test.

## Discussion

Although CRISPR-Cas9 technologies have immense promise in numerous aspects of biomedical science, many applications will benefit from tight temporal or spatial control over Cas9 activity, especially in the context of clinical development. Confining Cas9 activity to target cells and tissues of interest is highly desirable to prevent unforeseen adverse effects associated with off-tissue and off-target editing *in vivo*. Natural inhibitors of Cas proteins, anti-CRISPRs, can be repurposed as tools to limit the potential for unwanted edits. Acrs have several potential advantages for implementation as regulators. They are natural and genetically encodable inhibitors of Cas nucleases that have evolved as powerful inactivators of CRISPR immunity, usually offering some degree of specificity for particular types of systems. Moreover, their inhibition is often tunable/titratable based on the relative expression levels of Acrs and the target effectors, based upon stoichiometric mechanisms of action for most of them (van Gent and Gack 2018; Bondy-Denomy 2018). Most Acrs are small proteins that can tolerate fusions of fluorescent proteins or epitope tags, which could make them convenient for *in vivo* delivery by viral vectors or mRNAs and detection by fluorescence.

Here, we present a proof-of-concept demonstration of anti-CRISPR regulation by endogenous miRNAs *in vivo*, yielding tissue-specific control over CRISPR-Cas9 editing. We demonstrated that miRNA-mediated inhibition of anti-CRISPRs bearing hepatocyte-specific miR-122 MREs allows genome editing in a human hepatocyte cell line, Huh-7. Although this study used AcrIIC3_*𝒩me*_ for type II-C Nme1Cas9 and Nme2Cas9, as well as AcrIIC4_*Lmo*_ for SpyCas9, any well-validated combination of Acr-Cas nuclease will be compatible with this strategy, making it a versatile platform. With the wealth of new Acrs emerging for different CRISPR effectors (e.g. Cas12a; (Watters et al. 2018; Marino et al. 2018), we expect that opportunities for implementing this strategy will continue to increase. We also note that expression profiles of many miRNAs are well-defined for many tissues at many developmental stages and in numerous disease states (Alvarez-Garcia and Miska 2005). For example, miR-1 is highly and specifically expressed in cardiac and skeletal muscle tissues (Horak et al. 2016). The miRNA-repressible Acr system affords great flexibility in changing editing tissue specificity, given the ease with which the MREs can be swapped in the 3’UTR of the Acr transcript. Furthermore, because MREs are so small, this approach is well suited for viral modes of delivery (given the genome capacity constraints of viral vectors), and could confer specificity for some tissues that lack vector-compatible, tissue-specific promoters.

We extend this strategy to animal studies that document anti-CRISPR efficacy during Cas9-mediated editing *in vivo*. To our knowledge, this is the first demonstration of *in vivo* expression of Acr proteins in vertebrate models to inhibit Cas9 editing activity. From this study, we did not observe overt toxicity in the transfected liver tissues, although the safety and immunity profiles of delivered Acr proteins will need to be examined over longer periods of time and in additional biological contexts.

We exploited endogenous miRNAs for spatial control of anti-CRISPR expression to achieve tissue-specific editing by Cas9 *in vivo*. The endogenous miRNA repertoire has been combined with the CRISPR-Cas machinery previously to regulate the expression of Cas9 itself (Hirosawa et al. 2017; Senís et al. 2014). Whereas detargeting Cas9 expression from the liver (e.g. with miR-122) will allow editing to occur everywhere <u>except</u> the liver, our strategy will restrict Cas9 activity to the liver itself and protect all the other tissues. This will be particularly useful to restrict Cas9 genome editing to a single desired tissue following a systemic Cas9 delivery by AAV. Our results complement a strategy described by Wang *et al.*, which exploits miRNAs to release sgRNAs from longer, inactive precursors (Wang et al. 2019), though this approach has not yet been validated in tissue-specific editing applications *in vivo*. While this manuscript was in preparation, Hoffman *et al.* also reported using miRNA-regulated Acr proteins to achieve cell-type specific editing in hepatocytes and myocytes in culture (Hoffmann et al. 2019). Our studies further demonstrate that miRNA-repressible anti-CRISPRs can be applied in the tissues of adult mammals *in vivo*.

## Materials and Methods

### Vector construction

Codon-optimized AcrIIC3_*𝒩me*_ and AcrIIA4_*Lmo*_ sequences were ordered as gBlocks (IDT) and amplified using the primers with overhangs to the pCSDest vector by NEBuilder® HiFi DNA Assembly (NEB). Similarly, an mCherry ORF was fused to the C-terminus of each Acr by HiFi DNA assembly (NEB). To insert 3xmiR122 MREs in the 3’ UTR of each Acr, top and bottom strands were ordered as oligos (IDT) with restriction sites for SacI and HindIII and annealed before ligating into the vector linearized with the same restriction enzymes. For *in vivo* work, we used the hNme2Cas9-sgRNA_Rosa26 all-in-one AAV vector (Edraki et al. 2018). To make scAAV vectors expressing Acr proteins, the original scAAV plasmid encoding an EGFP ORF (a kind gift from J. Xie and G. Gao) and pCSDest-Acr plasmids were digested with SacI and AgeI restriction enzymes and then ligated. The sequences of codon-optimized Acr constructs and miRNA-122 MREs are also provided in the Supplemental Table 1. All plasmids used in this study are summarized in Supplemental Table 2 and will be available on Addgene.

### Cell culture and transfection

HEK293T and Huh-7 cell lines were cultured in Dulbecco’s modified Eagle’s medium supplemented with 10% fetal bovine serum (Sigma) and 1% penicillin-streptomycin (Gibco). For editing experiments *in vitro*, a total of 150 ng of Cas9, 150 ng of sgRNA, and 50 ng of Acr plasmids were transiently transfected in a 24-well format using Lipofectamine 2000 (Invitrogen) according to the manufacturer’s protocol. For Western blot analysis, 500 ng of each Acr vector and GFP plasmid used as a transfection control were transfected in a 6-well format using Lipofectamine 2000 (Invitrogen). The total DNA amount was kept constant by adding a stuffer plasmid in all cases.

### Flow cytometry

Transfected cells were trypsinized, washed in PBS, and resuspended in PBS for analysis on a MACSQuant® VYB from Miltenyi Biotec. A yellow laser (561 nm) with a 615/20 nm filter and a blue laser (488 nm) with a 525/50 nm filter were used for mCherry and GFP detection, respectively. Subsequent analysis was performed using FlowJo® v10.4.1. Cells were first sorted based on forward and side scattering (FSC-A vs SSC-A), and then single cells were gated using FSC-A and FSC-H. Finally, mCherry-positive cells were recorded after gating for GFP-positive (transfected) cells.

### Western blots

Proteins were collected 48 hours post-transfection and their concentrations were measured using the Pierce BCA Protein Assay Kit (Thermo Fisher Scientific). Western blots were performed as described previously (Lee et al. 2018) with primary mouse anti-FLAG (AbClonal, 1:5000) used for Acr detection and rabbit anti-HSP60 (1:5000) used for a loading control. After incubation with secondary anti-Rabbit or anti-Mouse antibodies (LI-COR IRDye®, 1:20,000), blots were visualized using a LI-COR imaging system.

### Mouse studies

C57BL/6 mice were obtained from Jackson Laboratory and all animal maintenance and procedures were performed following the guidelines of the Institutional Animal Care and Use Committee of the University of Massachusetts Medical School. Plasmids for hydrodynamic tail-vein injection were prepared using the EndoFreeMaxi kit (Qiagen). For hydrodynamic liver injection, a total of 90 ug of endotoxin-free plasmids was suspended in 2 ml of injection-grade saline and injected via the tail vein into 8- to 10-week-old C57BL/6 mice. Mice were euthanized 7 days post-injection and liver tissues were collected and stored at −80°C for analyses.

### Indel analysis

Genomic DNA from cells or liver tissues were collected using DNeasy Blood & Tissue Kit (Qiagen). Target sites were amplified using High Fidelity 2X PCR Master Mix (NEB). Primers used for PCR are listed in Supplementary Materials. PCR products were purified using DNA Clean & Concentrator Kit (Zymo) and sent for Sanger sequencing to obtain trace files (Genewiz). Indel values were estimated using the TIDE web tool (https://tide-calculator.nki.nl/).

### Statistical analysis

Standard deviations are derived from each group that has a minimum of three independent replicates unless otherwise noted. Unpaired, two-tailed t-test was used to determine the statistical significance between each group. Resulting P-values < 0.05, 0.01 and 0.001 are indicated by one, two, or three asterisks, respectively.

### Imunohistochemistry

Liver tissues were fixed in 4% formalin overnight, paraffin-embedded, and sectioned at the UMass Morphology Core. For Supplemental Figure 2, sectioned slides were stained with H&E for pathology analysis. For IHC, liver sections were dewaxed, rehydrated, and stained following standard protocols previously described (Xue et al. 2011) with primary antibodies against 3xHA-tagged Nme2Cas9 (anti-HA; Cell Signaling) and mCherry (anti-RFP; Rockland). Representative images are shown.

## Author Contributions

J.L. constructed all new plasmids used in this study, conducted all cell culture experiments, and analyzed samples derived from *in vivo* experiments. H.M. and S.Q.L. performed hydrodynamic injection and mouse tissue collection with guidance from W.X. R.I. provided tissue samples from mice injected with AAV8. J.L. and E.J.S. wrote the manuscript and all authors edited the manuscript.

## Supplemental Materials

Supplemental materials are available for this article.

## Acknowledgments

We are grateful to Jun Xie, Guangping Gao, and members of the Xue and Sontheimer labs for helpful discussions and sharing resources. We also thank Jordan Smith for assistance with IHC experiments, as well as Kevin Luk, Pengpeng Liu, and Scot Wolfe for sharing sgRNA plasmids. This work was supported by grants from the U.S. National Institutes of Health (GM125797) to E.J.S and (DP2HL137167 and UG3HL147367) to W.X. as well as institutional funds to W.X. and E.J.S.

---

**Supplemental Figure 1.**
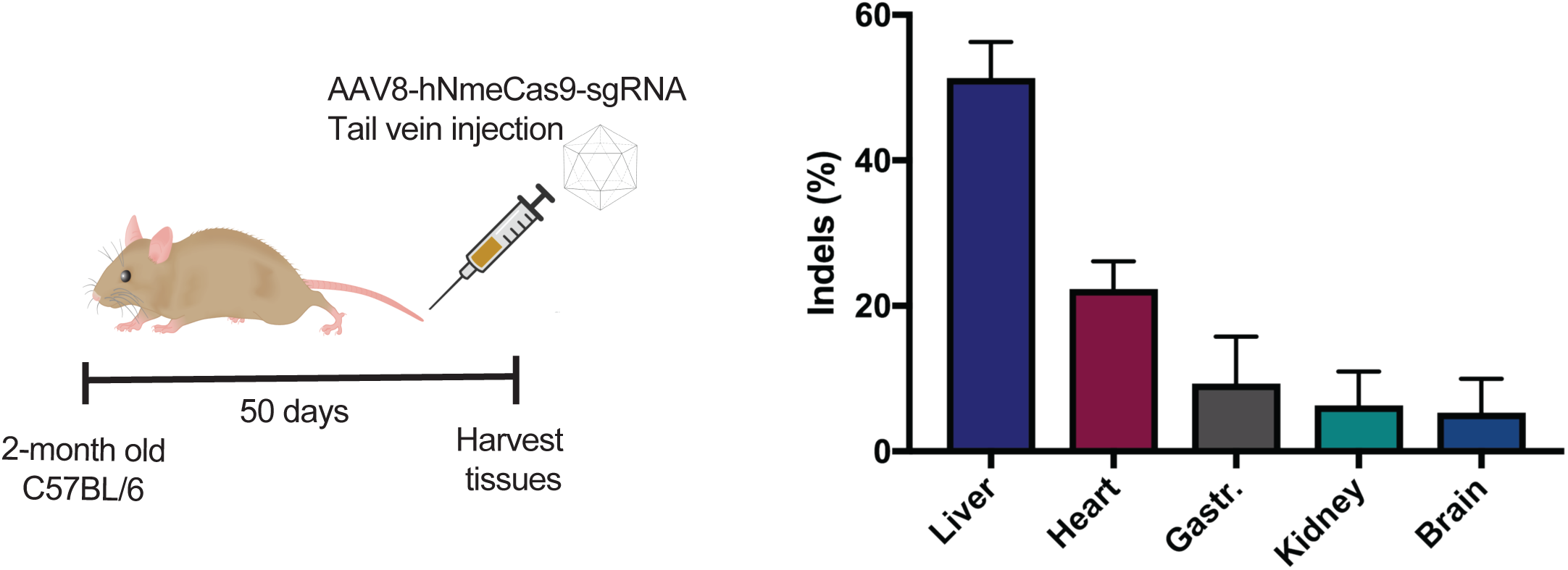
Editing in different organs collected 50 days after rAAV8 delivery of all-in-one hNme1Cas9/sgRNA targeting *Rosa26* via tail vein injection in C56BL/6 mice (n = 3). Indels are measured by TIDE analysis. Gastr., gastrocnemius muscle.

**Supplemental Figure 2.**
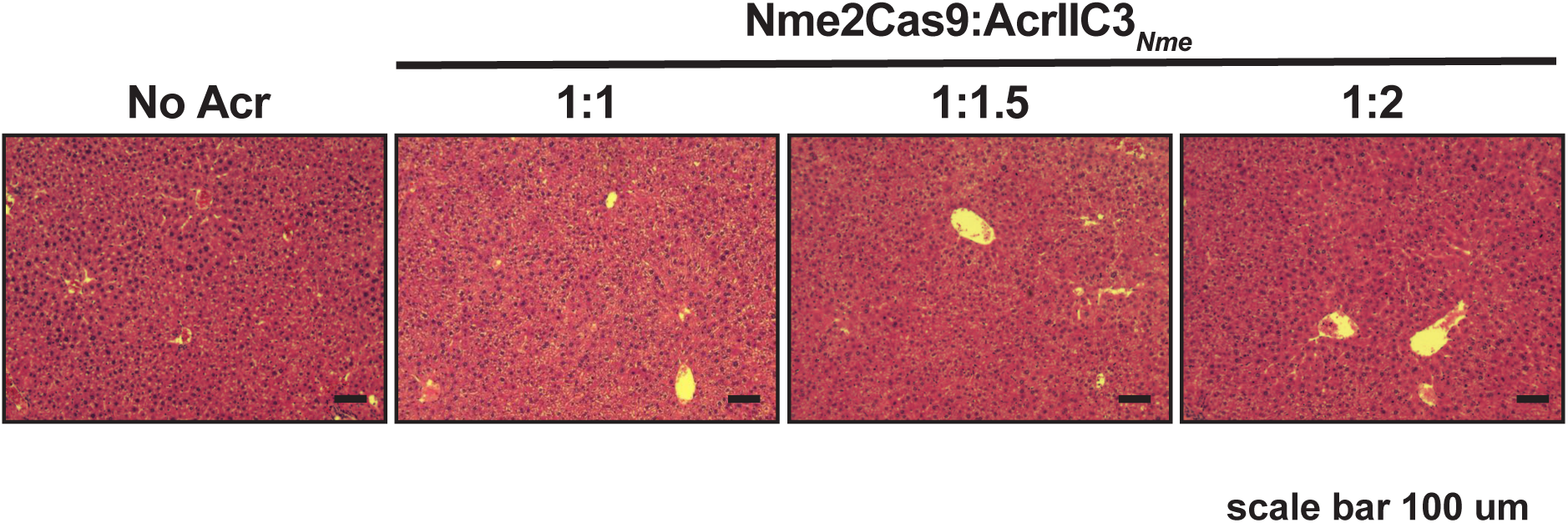
H&E staining of liver tissue sections from mice injected with hNme2Cas9 and AcrIIC3_*𝒩me*_ expression plasmids at different ratios exhibit no overt toxicity. Scale bar, 100 µm.

**Supplemental Figure 3.**
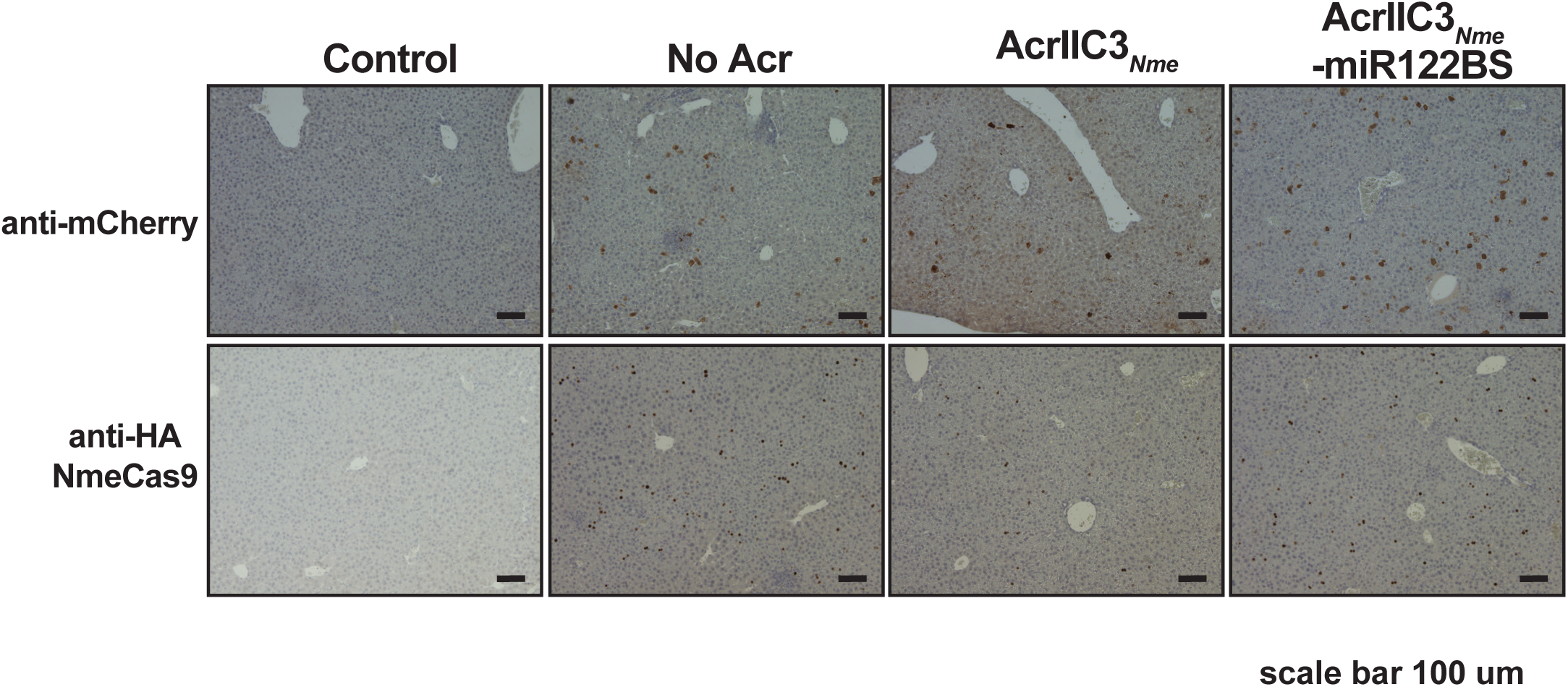
Immunohistochemistry of liver tissues from mice injected with hNme2Cas9/sgRNA plasmid alone, with AcrIIC3_*𝒩me*_ plasmid, or with AcrIIC3_*𝒩me*_-3xmiR122BS, as in Fig. 4D. Anti-mCherry was used to detect mCherry expression from injection control plasmids. Anti-HA was used for 3xHA tagged hNme2Cas9 detection. Control, saline-injected. Scale bar, 100 µm.

## Supplemental Materials

Table 1. Sequences of codon-optimized anti-CRISPR proteins.

Table 2. Plasmids and oligonucleotides used in this study.

---

**Supplemental Table 1.**
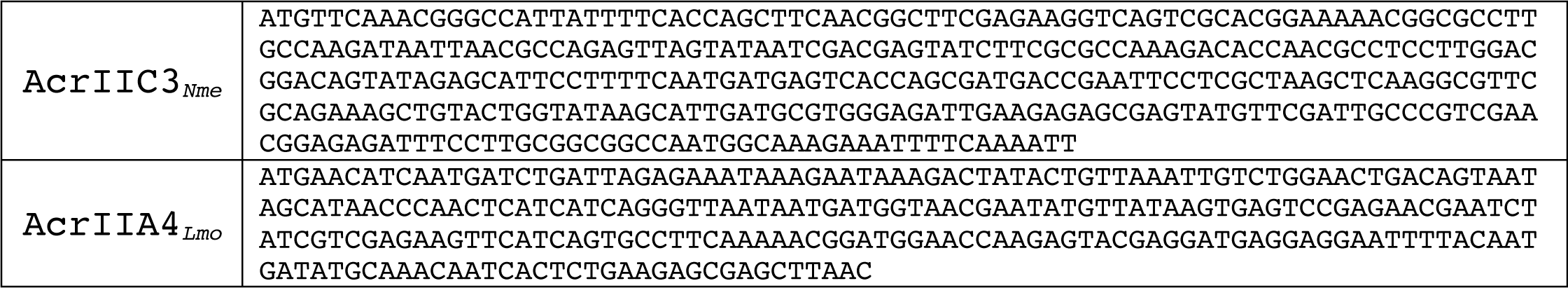
Sequences of codon-optimized anti-CRISPR proteins.

**Supplemental Table 2.**
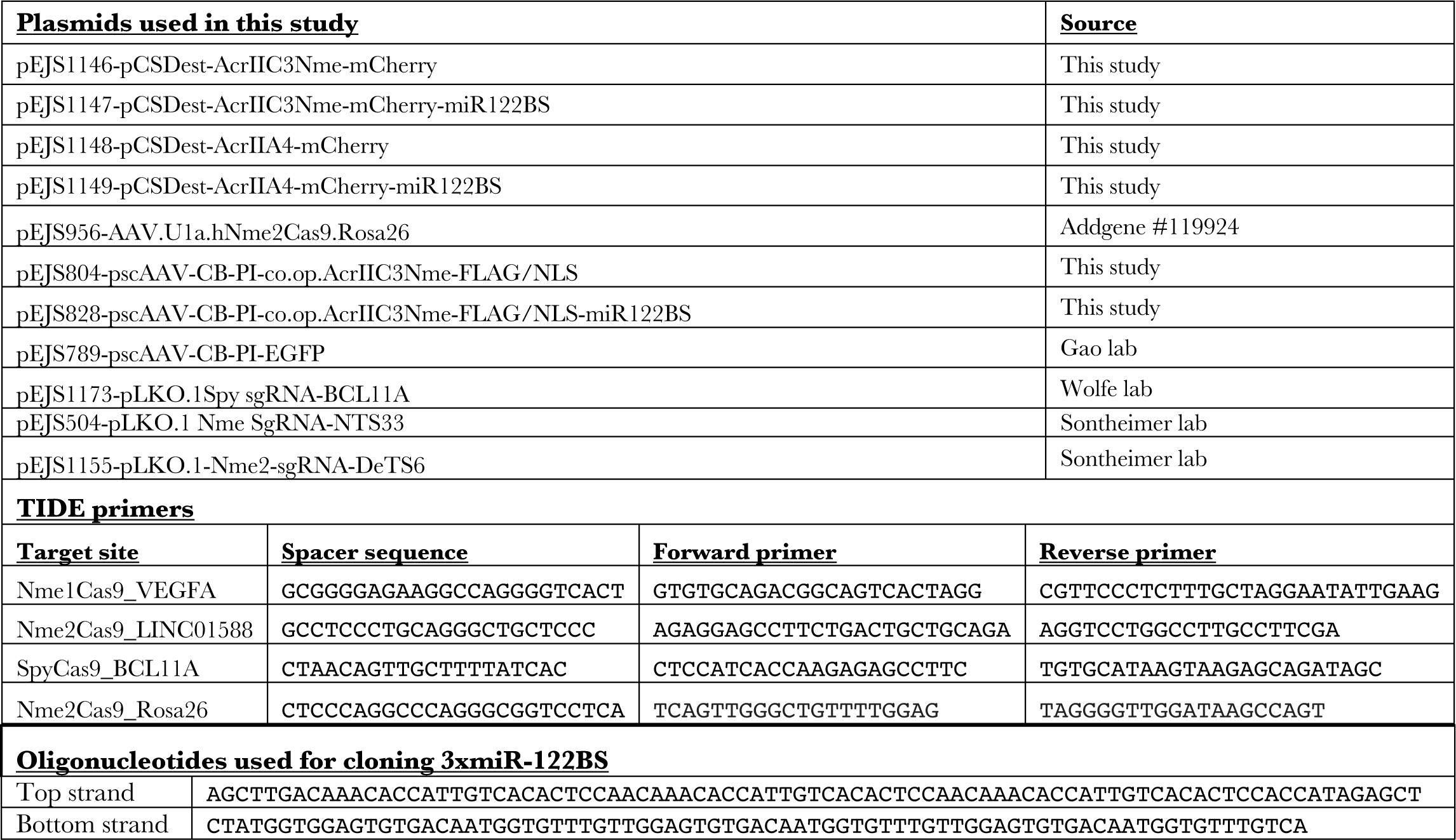
Plasmids and oligonucleotides used in this study.

